# Automated long-term two-photon imaging in head-fixed walking *Drosophila*

**DOI:** 10.1101/2021.03.20.436241

**Authors:** Andres Flores Valle, Rolf Honnef, Johannes D. Seelig

**Affiliations:** Center of Advanced European Studies and Research (caesar), Bonn, Germany

## Abstract

The brain of *Drosophila* shows dynamics at multiple timescales, from the millisecond range of fast voltage or calcium transients to functional and structural changes occurring over multiple days. To relate such dynamics to behavior requires monitoring neural circuits across these multiple timescales in behaving animals.

Here, we develop a technique for automated long-term two-photon imaging in fruit flies, during wakefulness and sleep, navigating in virtual reality over up to seven days. The method is enabled by laser surgery, a microrobotic arm for controlling forceps for dissection assistance, an automated feeding robot, as well as volumetric, simultaneous multiplane imaging. The approach is validated in the fly’s head direction system.

Imaging in behaving flies over multiple timescales will be useful for understanding circadian activity, learning and long-term memory, or sleep.

## Introduction

The brain of *Drosophila* shows dynamics at multiple timescales, from single action potentials to functional and structural changes across multiple days. Such slow changes occur for example due to sleep^1^, circadian rhythms^2, 3^, or memory consolidation^4, 5^. To relate fast and slow activity changes across these conditions therefore requires monitoring neural circuits over multiple timescales in behaving animals^6^.

While head-fixed walking preparations allow imaging of neural activity during behavior^7^, experiments typically last on the order of an hour, and two-photon imaging^8^ over multiple timescales, including naturally occurring sleep, has so far not been achieved. A preparation for repeated imaging over up to 50 days through a transparent window allowed functional imaging in an epifluorescence widefield microscope after anesthetizing and reintroducing the fly into the setup for each experiment^9^. Three-photon imaging of neural activity in intact walking flies over 12 hours was performed for bright and large neurons close to the surface of the brain^10^.

Here, we describe a method for multi-day two-photon calcium imaging in behaving fruit flies over up to seven days. Continuous monitoring of behavior, which included sleep and circadian modulation, in a virtual reality setup was combined with intermittent, fast volumetric imaging of calcium activity during between 8 to 16 percent of the time (imaging every 5 minutes in trials of either 30 or 60 seconds throughout the duration of the experiments).

For this, a transparent window was inserted into the fly’s head^9^, using a cost effective laser surgery approach combined with a microrobotic arm for operating forceps under a dissection microscope. The imaging and behavior experiments did not require supervision, with an automated feeding system maintaining the fly over the course of the experiment. Simultaneous two-plane imaging^11^ allowed volumetric recording of neural activity at high frame rates. We validated the method in wedge neurons in the head direction system of the fly^12, 13^.

## Results

### Laser surgery

A window for imaging through the cuticle was cut using a laser^9^ (Fig. 1a and b, Supplementary Video 1, and Methods). Different from previous approaches^9, 14^, which used an expanded pulsed UV excimer laser at 193 nm, we here used a focused visible continuous wave laser (Lassos Lasertechnik, YLK Series, 561 nm) with comparatively low power (30 mW was required for cutting) and cost. The fly (glued to a pin with its head fixed with respect to the thorax, see Methods) was moved against the stationary laser focus using a cost-effective, three-axis motorized micromanipulator (see Methods 1.1). A path for the laser was defined in three dimensions on the fly’s head, allowing arbitrary geometries for the resulting opening in the cuticle (Fig. 1b). A single pass through the laser focus was sufficient to cut the cuticle.

**Figure 1.**
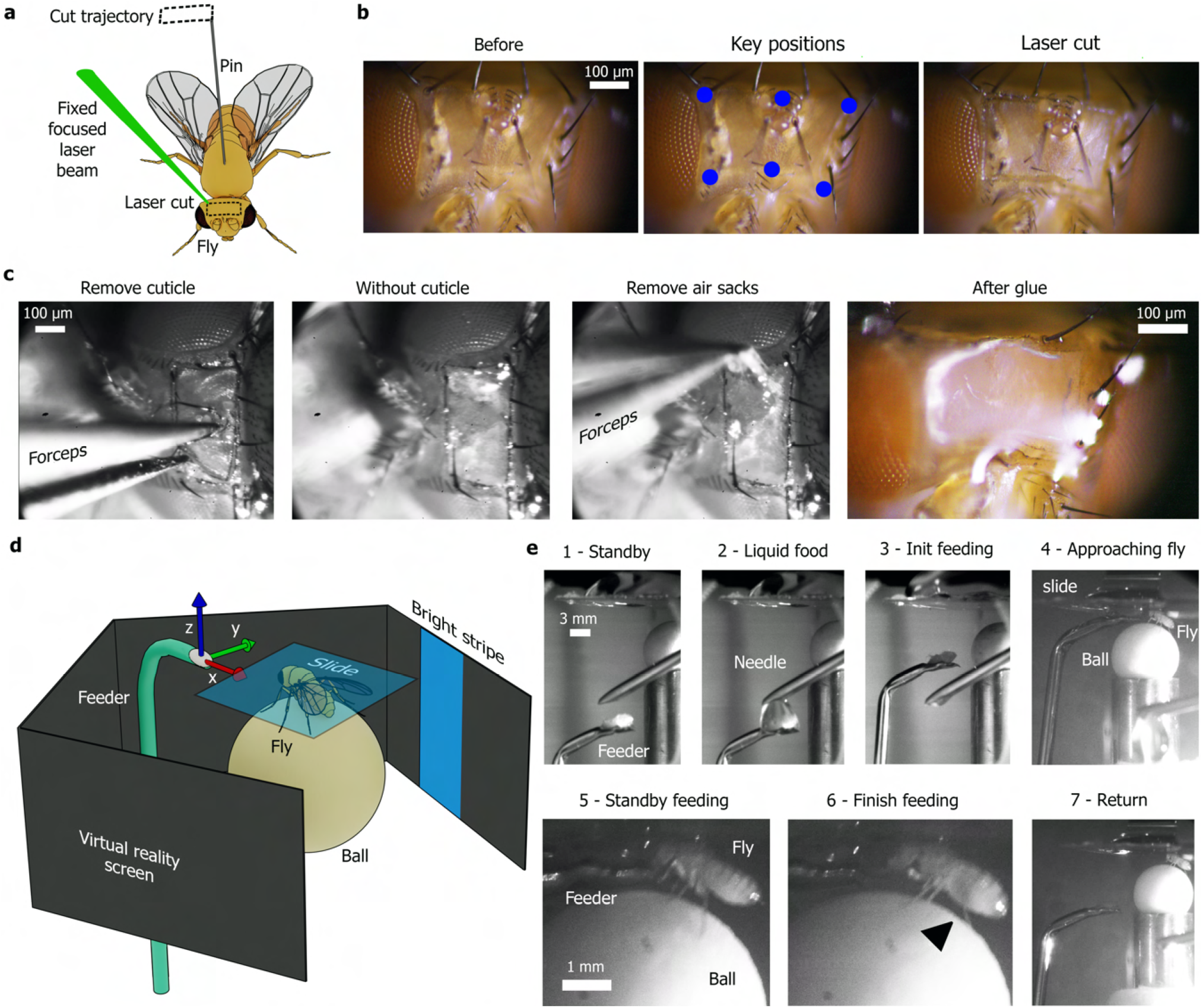
Long-term imaging preparation. **a** Schematic of laser cutting. The fly attached to a pin is moved while a laser beam cuts the cuticle. **b** Head of the fly (top view) before laser cutting, with key positions of trajectory, and after cutting. **c** Top view of head with motorized forceps removing cuticle and air sacs. The window is sealed using transparent glue^9^. **d** Schematic of VR and feeding setup: the fly is glued to a slide and placed on a ball under the microscope (not shown) surrounded by a screen for virtual reality projection and an automated feeding system. **e** 1 - Side view of fly and feeder in standby position (4 hours). 2 - Refilling of feeder every 2 minutes. 3, 4 - Feeder approaches fly. 5 - Feeding the fly (2 minutes). 6 - Increased abdomen volume from the ingested food indicated with arrowhead. 7 - Feeder returns to home position.

### Microrobotic arm for surgery assistance

After laser surgery, the fly on the pin was transferred to a dissection microscope. The pin was attached to a micromanipulator and was positioned on a cold plate at 4 degrees Celsius to anesthetize the fly. To assist with dissections, a microrobotic arm for controlling forceps was used to lift off the cut cuticle as well as to remove the underlying air sacs (see Methods 1.3, Fig. 1c and Supplementary Video 2). The position, orientation, and grip of the pair of forceps were controlled with a joystick.

The forceps were positioned at an angle of 45 degrees with respect the table and the fly was positioned on a cold plate angled downwards at approximately 20 degrees with respect to the table to easily access the brain. The tip of the forceps (see Fig. 2 and Methods 1.3 for details) was inserted into the cut to grab and slowly retract the cuticle (Supplementary Video 2). After cleaning the tip of the forceps, it was repositioned and the air sacs were similarly removed (Supplementary Video 2).

**Figure 2.**
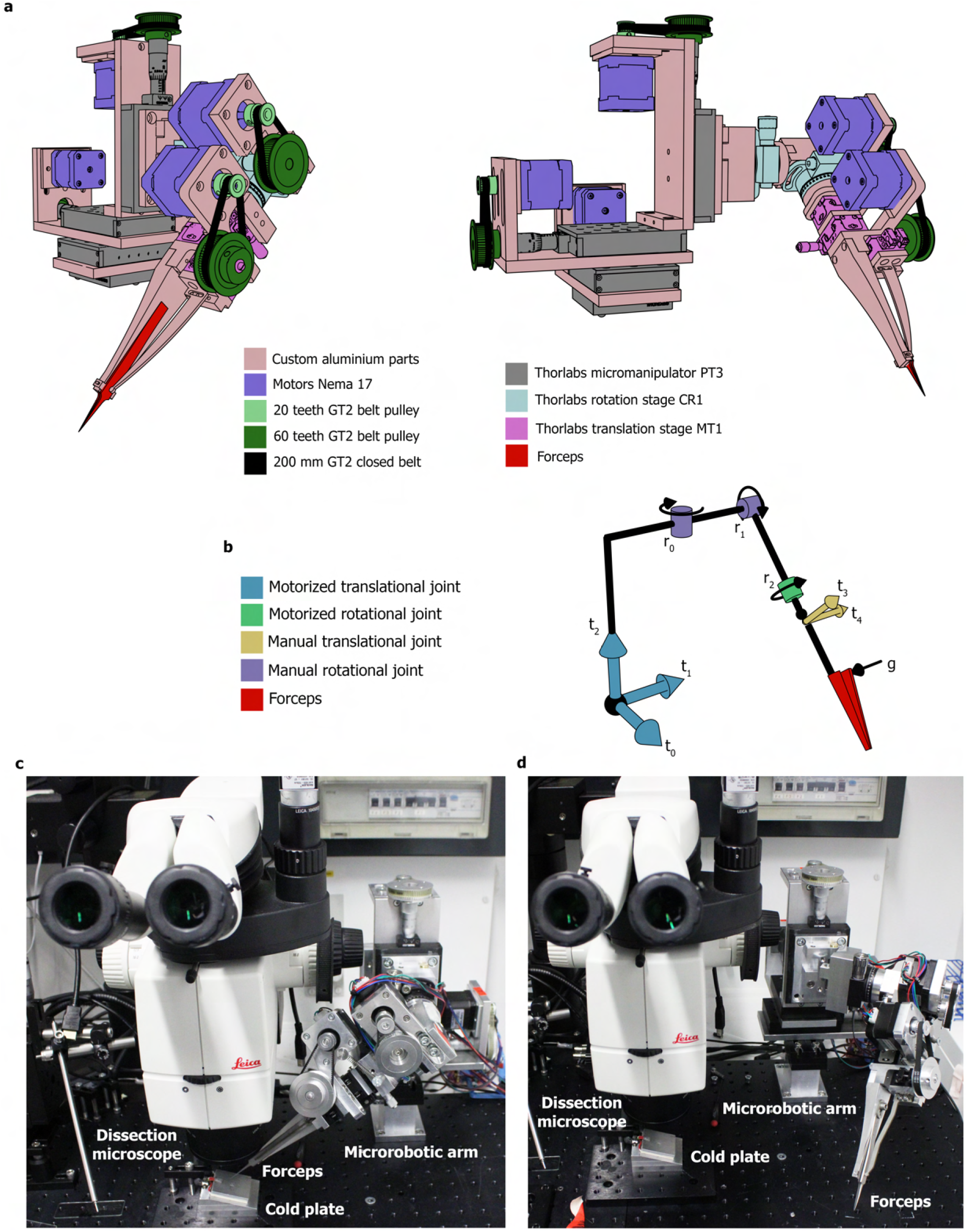
Microrobotic arm for dissection assistance. **a** Two different views of 3D model of microrobotic arm with forceps. **b** Scheme of dynamics with different joints and forceps. **c** Setup with microrobotic arm with forceps under the dissection microscope. **d** View of setup with retracted forceps (using joint *r*_0_), for example for cleaning the tip of the forceps during dissections.

The opening in the fly’s head was sealed by manually applying UV glue^9^ with a pin, here we used UV highgloss finish (DETAX, Freeform, 02204), and cured for twice 15 seconds with UV light. The fly was then glued to a microscope cover slide for imaging, either after first leaving the fly to recover overnight in a vial with food^9^ (fly 3, Supplementary Fig. S6 and S7, Supplementary Video 6, and fly 4, Supplementary Fig. S8 and Supplementary Video 7) or directly after surgery (fly 1, Fig. 4 and Supplementary Video 4, and fly 2, Supplementary Fig. S5 and Supplementary Video 5). The fly glued to the microscope slide was finally positioned on an air supported ball in a VR system (Fig. 1d) under a two-photon microscope^7, 15^.

### Volumetric two-photon imaging

For two-photon imaging of calcium activity we implemented a fast volumetric approach with two axially extended Gaussian beams. The focal planes of the two beams were axially offset and recorded simultaneously using temporal multiplexing^11, 16, 17^ (Fig. 3b) which allowed covering the entire structure of interest in a single scan at 60 Hz. The extended imaging volume also minimized artefacts resulting from axial brain motion and corresponding changes in fluorescence intensity.

**Figure 3.**
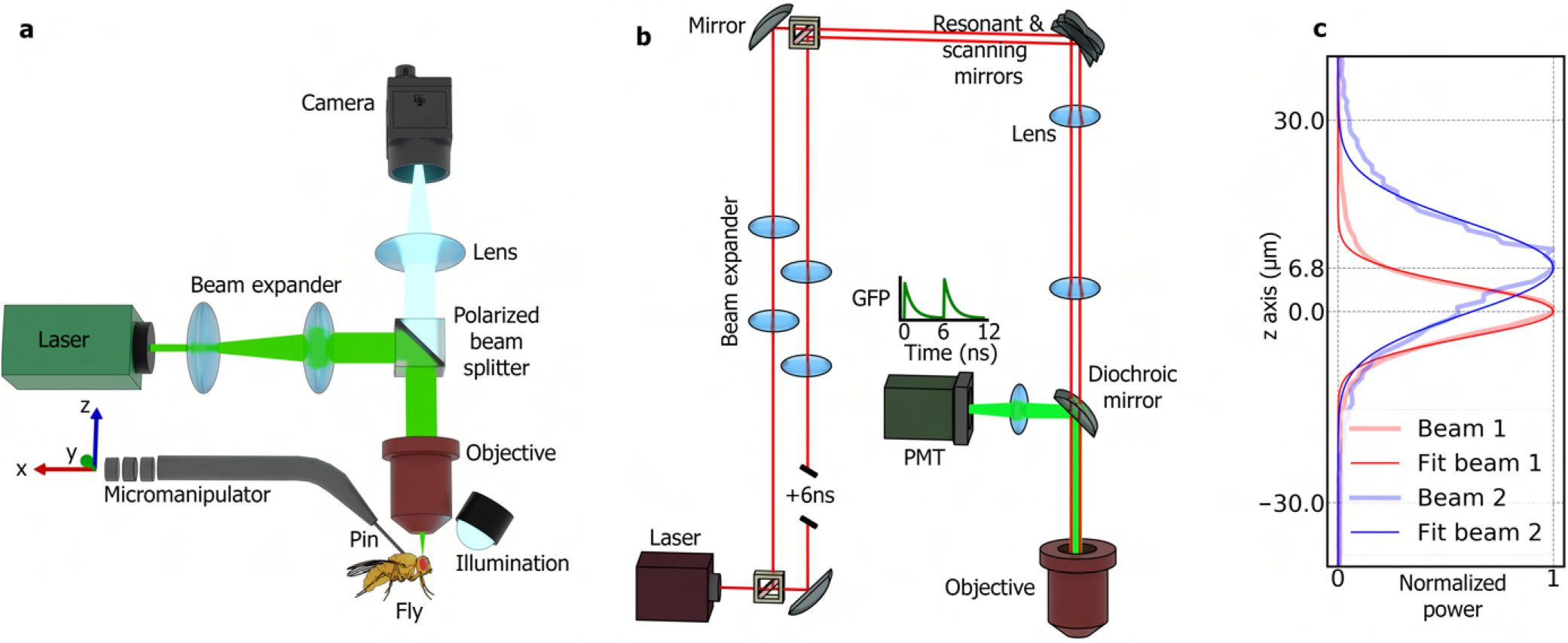
Optical setups. **a** Schematics of setup for laser cutting. The laser beam is fixed and the fly is moved along three axes (*x, y* and *z*) with a custom motorized micromanipulator. A camera and custom microscope are used to set the key points on the fly’s head that define the cutting trajectory. **b** Two-photon microscope with two Gaussian beams that are temporally offset by 6 ns for simultaneous imaging in two different focal planes using temporal multiplexing^11^. **c** Normalized axial (*z* axis) profiles of the two beams fitted with a Gaussian function.

The laser power at the sample was 6 mW for each beam. The microscope optics and electronics were similar to the one described in^17^ with temporal mutliplexing implemented in the Scanimage photon counting mode^17, 18^. The two beams had a delay of 2 m and were linearly and orthogonally polarized (Fig. 3b). Beam diameter and collimation were adjusted with two lenses (Thorlabs achromatic doublets). The microscope objective (16X Nikon CFI LWD Plan Fluorite Objective, 0.80 NA, 3.0 mm WD) was underfilled, resulting in elongated beam profiles with an axial standard deviation of 4*μ*m and 7*μ*m, respectively, and the beam maxima were offset by 7*μ*m (Fig. 3c).

For long-term imaging experiments, data was recorded in continuous trials of 30 (flies 3 and 4) or 60 seconds (flies 1 and 2), with a break of 5 minutes between trials. Recordings were stopped for 2 minutes per day for refilling the liquid food deposit. The number of trials, the time of each trial, the total time during which calcium activity was recorded, as well as the total time and temperature of the experiment are shown for each fly in Supplementary Table S1.

### Feeding robot

To maintain the fly over multiple days in the VR setup, a robot was developed for automated feeding of the fly on the ball (Fig. 1d and e). A feeder tube can be used for imaging the midgut over up to 16 hours^19^. Installing such a tube during the entire duration of the experiment was however not compatible with virtual reality, where it blocks the fly’s view. Therefore a motorized micromanipulator (see Methods 1.1) was used to automatically introduce and remove a feeder at preset intervals during the experiment. The feeder consisted of an appropriately shaped needle (Fig. 1e and Supplementary Fig. S2b) with a piece of cotton inserted into the tip^19^. This needle was parked in a standby position outside the fly’s field of view and only approached the fly for feeding (Fig. 1d, e, Supplementary Fig. S2b and Supplementary Video 3). At the beginning of a multiday experiment, the feeding position of the needle was defined such that the fly could extend its proboscis into the food (as monitored with two cameras (Basler acA640-750um) from different angles, Fig. 1 d). Further, a trajectory for approaching the needle to the fly’s proboscis and then retracting it to a standby position outside the fly’s field of view was defined (Fig. S2 b and Supplementary Video 3). A second needle continually refilled the feeding needle in the standby position with liquid food to keep the cotton moist. This refilling needle was connected through tubes to an elevated liquid food container with an electromagnetic valve (electric solenoid valve working at 24V) for timing the refilling process (see Figure S2b). The valve was controlled by an Arduino Uno connected to a host computer that opened the valve every 2 minutes for 0.5 seconds.

The feeder was programmed to feed the fly for 2 minutes every 4 hours. The intervals and duration for feeding were chosen such as to maintain the fly over multiple days but also to ensure sufficient walking activity (which typically decreased after feeding) for monitoring activity in the head direction system of the walking fly. For fly 1, the feeding process was interrupted during the second night and required an extra feeding epoch outside of the scheduled times (Fig. 4, top row and 11th red vertical line) which resulted in the increased walking activity during the night.

**Figure 4.**
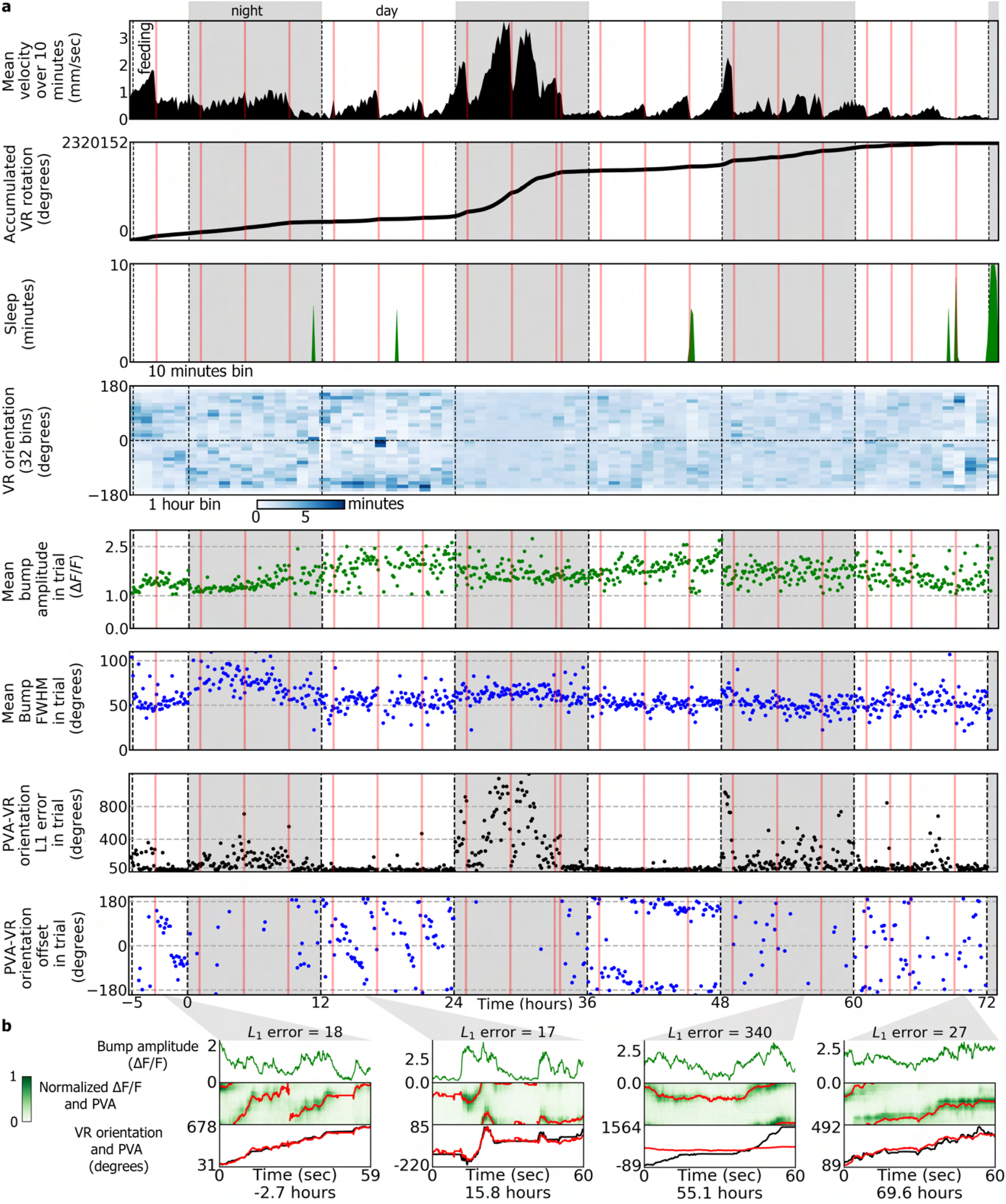
Imaging and behavior experiment for fly 1. **a** First row: velocity of the fly averaged in bins of 10 minutes. Vertical red lines indicate feeding, while white and grey regions indicate day and night. Second row: accumulated rotation of the fly in the VR. Third row: sleep time in bins of 10 minutes. Fourth row: time distribution of the VR orientation. The VR orientation is divided into 32 angular bins and 1 hour temporal bins. Fifth row: mean bump amplitude of wedge neurons in each trial. Sixth row: mean bump FWHM of wedge neurons in each trial. Seventh row: L_1_ error between the PVA and the VR orientation in each trial. Last row: offset between the PVA and VR orientation in each trial. **b** Example trajectories of activity in wedge neurons in different trials at different times in the experiment. For each of the four trials, top row: amplitude of the bump. Second row: change in fluorescence of wedge neurons (green image) together with the calculated PVA (red line). Bottom row: unwrapped trajectory of the VR orientation (black line) together with the PVA (red line) with PVA-VR orientation offset removed. L_1_ error is indicated at the top for each trial.

### Calcium imaging over multiple days

For validation of the long-term imaging approach, we monitored calcium activity in wedge neurons in the head direction system of flies walking with a single bright stripe in a VR setup^12^ during the day (12 hours) and in darkness during the night (12 hours). The temperature of the preparation was controlled with a perfusion system circulating water under the objective (see Methods 1.5 and Supplementary Fig. S2c). Calcium activity was recorded in trials of 60 (flies 1 and 2) or 30 seconds (flies 3 and 4) every five minutes over a total duration of up to seven days, while behavior was monitored continuously.

Wedge neurons encode the head direction of the fly, for example with respect to a bright stripe, in a bump of activity that moves along the ellipsoid body (EB)^12^. To show that neural activity could be recorded reliably in walking animals over multiple days, we therefore verified that the bright stripe in the VR was accurately tracked by the bump in the EB. Bump position (measured with the population vector average (PVA) over 32 regions of interest tiling the EB, Supplementary Fig. S4b) tracked the bright bar during the day (as measured with *L*_1_ error between the population vector average of bump activity and the stripe position (equation 8)) and with an error during the night, as previously described^12^ (Figures 4 and 5, and Supplementary Figures S5, S6, S7, S8). This, together with the animal’s walking activity (see Supplementary Videos 4, 5, 6 and 7), indicated that neural activity and behavior could be recorded reliably.

**Figure 5.**
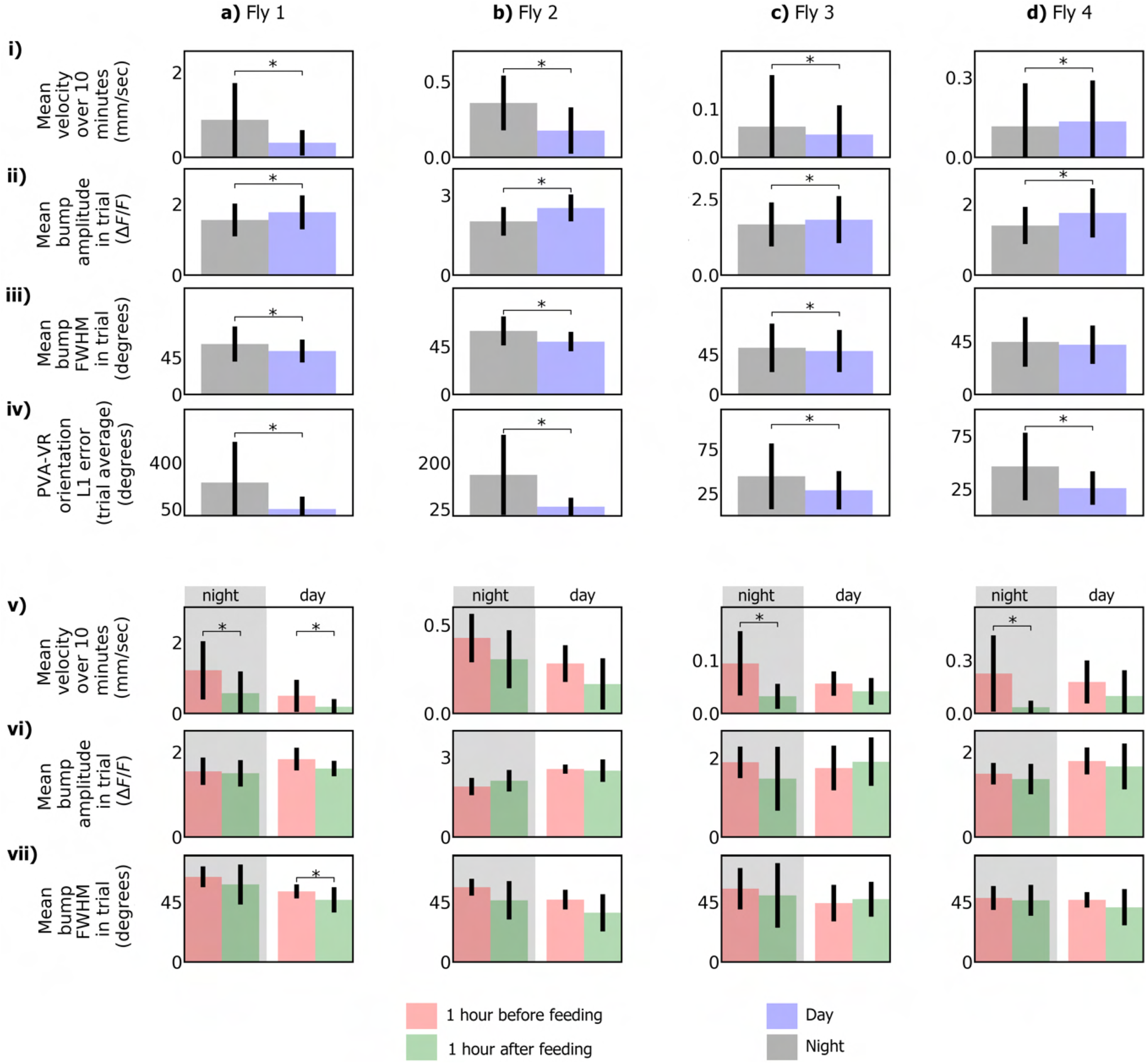
Statistical analysis of experiments. Top, row **i)**: mean velocity during day and night for fly 1 (**a**), fly 2 (**b**), fly 3 (**c**), and fly 4 (**d**). Row **ii)**: mean bump amplitude, row **iii)**: mean bump FWHM, and row **iv)** difference between bump position (PVA) and visual stimulus orientation (VR). Bottom, row **v)**: mean velocity during 1 hour before feeding and 1 hour after feeding for fly 1 (**a**), fly 2 (**b**), fly 3 (**c**), and fly 4 (**d**). Row **vi)**: mean bump amplitude. Row **vii)**: mean bump FWHM. Asterisks indicate statistical significance (p < 0.05).

Wedge neuron activity was typically higher during the day with visual stimulation than during the night (Fig. 5ii). Feeding did not impact wedge neuron activity, but did change walking activity (Fig. 5i). As observed in freely moving flies, walking activity was interrupted by bouts of sleep, both, during the day and night (Figures 4, 5, and Supplementary Figures S5, S6, S7, and S8) and (in fly 3) showed circadian modulation (Supplementary Figures S6 and S7). Sleep or walking activity were also not affected by the onset of the imaging trial and the switching on of the laser (Supplementary Video 8).

The offset between the bump and the stripe position, which was previously found to vary between trials and flies^12^, showed slow unidirectional drift of varying amount that changed between different days (see for example fly 1, Fig. 4, row 8 during days 2 and 3, respectively) independent of the amount of rotational walking activity. Such drift in bump offset could at least partially originate from occasional delayed movement of the bump with respect to the VR stimulus, which was observed in a few trials in fly 1 during the second day where PVA-VR orientation drift was high (Supplementary Fig. S10), but not during the first or third day. In five trials during the second day (Supplementary Fig. S10b and c), the bump was delayed around one of the wedges with respect to the VR, increasing the offset between the PVA and VR orientation before resulting in a jump (similar to discontinuous movements previously described in experiments with multiple visual stimuli^12^). A potential explanation for such delayed movement could be heterogeneity in the connectivity between wedge neurons^20^.

For flies that were used for imaging immediately after surgery (flies 1 and 2), the L_1_ error was large during the first hours of the experiment, even though a bump was visible, and this data was not included in the average statistics (Fig. 5). Changes in the dynamics over long timescales were also observed in fly 3 after 5 days. The L_1_ error increased while the correlations stayed high (Supplementary Fig. S9a), corresponding to a change in the gain of the head direction system with respect to ball rotation. Under these conditions, we observed (after the 5th day) autonomous activity of the bump while the fly was standing still on the ball during the night (Supplementary Fig. S9b). This likely indicates a deterioration of the head direction system, potentially due to phototoxicity, since overall walking behavior was not visibly affected.

## Discussion

We developed a preparation for functional calcium imaging over up to 7 days during navigation in virtual reality. The behavior and imaging experiments, using volumetric, simultaneous two-plane two-photon imaging, was fully automated and didn’t require intervention while the fly was maintained in the VR with an automated feeding system. The preparation opens up the opportunity to investigate neural circuits underlying behavior across multiple timescales, from fast calcium dynamics to changes that occur over multiple days. Monitoring behavior continuously, also between the intermittent imaging sessions as was done here, will be useful for relating behavior to neural activity changes in the context of circadian activity, learning and long-term memory, or sleep.

Functional imaging during sleep has so far only been performed in flies that were placed on the ball for short periods of time^21^. Similarly, circadian activity has so far only been investigated in immobilized animals^2, 3^. The long duration of our experiments resulted in the observation of multiple epochs of extended naturally occurring sleep. Fly walking activity depended on temperature and feeding (decreasing after feeding, Fig. 5), which could be adjusted to increase or decrease the amount of sleep or walking activity. The automated feeding setup will also be useful for studying feeding behavior or drug application in behaving animals during functional imaging.

The reproducibility and ease of dissections was improved compared to manual approaches^7^, only requiring minimal manual manipulations under the dissection microscope for applying glue. This, together with the extended lifetime and improved health of the fly, even without feeding, makes this approach also attractive for imaging experiments over short timescales (after leaving the flies to recover for several hours^9^). Compared to previous laser surgery approaches^9^, a comparatively low cost continuous wave laser was used for surgery in a scanning configuration. This approach allows greater flexibility in terms of head orientation and the geometry of the cut, which is defined in all three spatial dimensions and therefore is not limited to a single focal plane.

Compared to three-photon experiments in intact animals, which still need to be tested for calcium imaging in deeper structures, the approach used here has the advantage that it is compatible with more widely available one-or two-photon imaging approaches^9^. While previous approaches used widefield imaging for monitoring calcium dynamics^9^, increased fluorescence background makes this less suitable for imaging in deeper structures and visible excitation light interferes with behavior. We here therefore used two-photon imaging with an axially extended focal volume based on two simultaneously recorded focal planes using temporal multiplexing. These experiment could also be performed without multiplexing by averaging the signal of the two beams in a single detection channel.

Overall, long-term imaging in fruit flies walking in VR is expected to contribute to an understanding of the dynamics of neural circuits in a variety of behaviors that bridge multiple timescales, for example circadian rhythms, learning and long-term memory, or sleep.

## Acknowledgements

We would like to thank Zohaib Amir, Bernd Scheiding, the caesar mechanical workshop for help with electronic and mechanical components of the setup, Tim Krause and Mina Baayer for fly maintenance and fly food, and Ivan Vishniakou for comments on the manuscript.

## Author Contribution

AFV and JDS designed the experiment and wrote the paper. AFV performed all experiments and data analysis. AFV built all setups, JDS set up the microscope. AFV and RH developed the microrobotic arm.

## 1 Methods

### 1.1 Motorized micromanipulators for fly feeding and laser surgery

For both, feeding the fly on the ball as well as for moving the fly across the laser beam for surgery, we motorized a three-axis micromanipulator (Thorlabs, PT3 with 25 mm travel). Fig. S1a shows the mechanical assembly and the control configuration. Each axis was controlled with a stepper motor (Nema 17). Motor torque was transmitted to the micromanipulator actuator through a closed loop toothed belt (GT2 with 6mm width and 200mm length, black parts in Fig. S1a), that connected a metal gear attached to the motor axis (20 teeth GT2 belt pulley with 5mm bore, green parts in Fig. S1a) and a custom 3D printed gear with 92 teeth attached to the micromanipulator axis. The three motors were supported by 3D printed parts, shown in red in Fig. S1a. All 3D printed parts were printed using a Prusa, i3 MK3S printer. Using micro-stepping of the stepper motors (3200 steps per revolution) and gear reduction, the micromanipulator produced a translational resolution of 0.06*μ*m per motor step in each axis.

**Figure S1.**
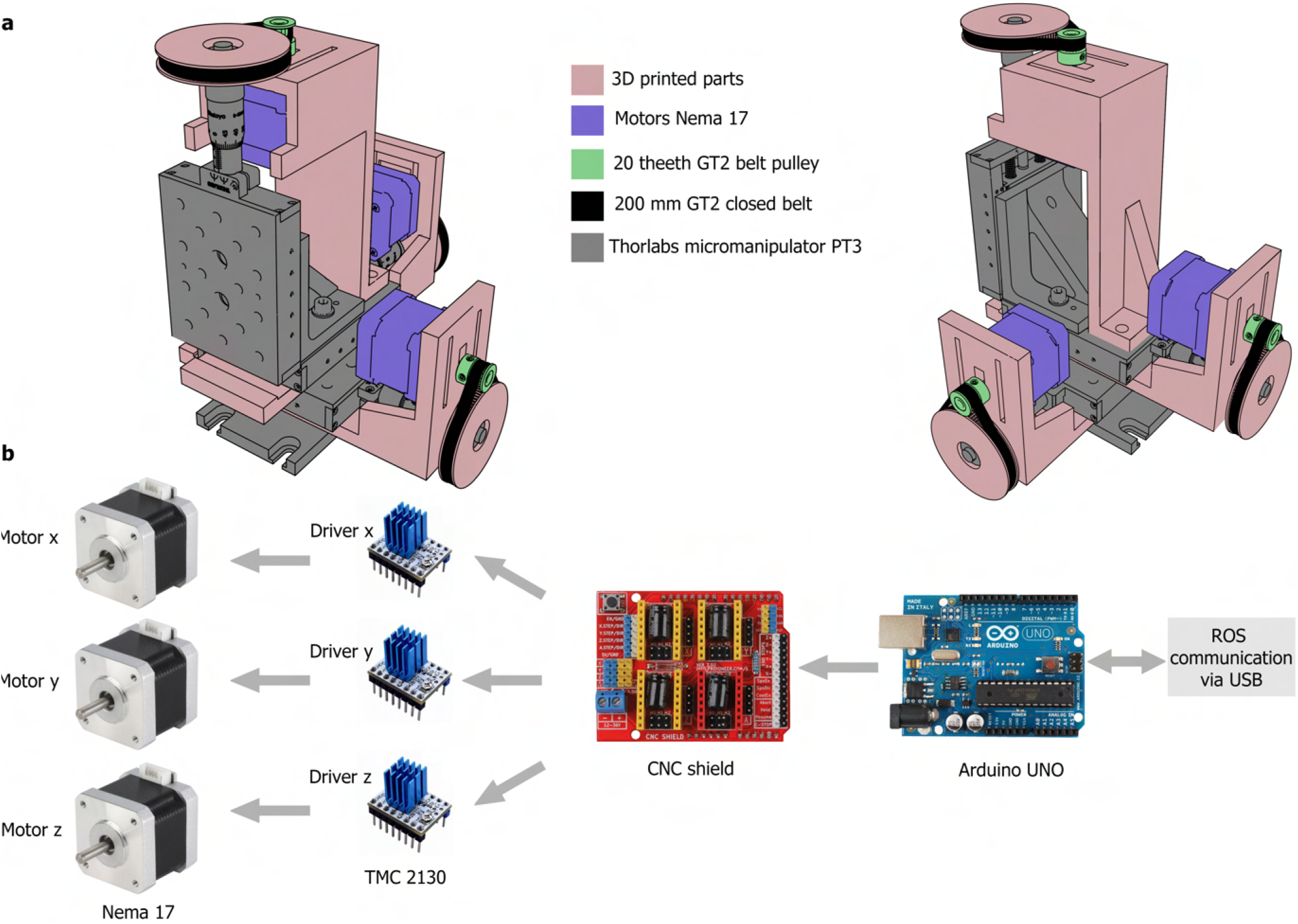
Motorization of micromanipulator. **a** Two views of 3D model with different parts labeled in different colors. **b** Electronics for controlling Nema 17 motors. Motors are controlled by TMC 2130 drivers attached to CNC shield connected to Arduino UNO. Arduino UNO is connected to a host computer that interacts within the ROS framework by receiving motor commands and sending motor absolute positions.

The stepper motors were controlled through Arduino Uno using a CNC shield with TMC2130 motor drivers powered with 12V. Arduino Uno was connected through USB to a host computer running ROS (Robot Operating System) under Linux. The connection diagram is shown in Fig. S1b. Custom control software implemented in Python sent motor commands through serial communication at 30 Hz, while Arduino Uno returns the absolute angular position of each motor at 10 Hz. Multiple micromanipulators could be connected to a single computer through USB ports.

Each micromanipulator was controlled in open loop through either velocity or absolute position commands from the host computer. A custom command line interface was developed in Python to control each micromanipulator with three control modes: (1) ’stop’, where the micromanipulator is standing still. (2) ’manual’, where the user can move the micromanipulator using a joystick (Logitech Extreme 3D pro joystick); velocity commands are sent to the motors proportional to the displacements of the joystick axes (absolute maximum velocity can also be adjusted from one of the joystick’s axes). (3) ’auto’ mode, in which the user can program instructions, via the command line interface, that the micromanipulator repeats sequentially in a loop. Additional instructions can be defined, such as a wait instruction, where the manipulator waits for a defined amount of seconds before moving to the next instruction. This is used in the feeding robot during the standby and feeding states (Fig. 1e, panels 1 and 5).

### 1.2 Laser surgery setup

The laser was expanded and collimated with two lenses to fill the back aperture of the microscope objective (Olympus Plan N 10x/0.25 air objective) over a dichroic mirror (Semrock). A white light LED was used to illuminate the fly and to image its head through the same objective with an additional lens (Thorlabs AC254-250-A) onto a color camera (Basler acA1920-155uc). Laser power was adjusted to 32 mW at the focal plane. The laser beam was additionally pulsed with a mechanical shutter (Thorlabs SH05/M, opened for 15 milliseconds every 45 milliseconds), resulting in an average power of 8 mW at the focus.

The fly, tethered to a pin, was attached to a motorized micromanipulator. This custom-motorized micromanipulator, based on a manual translation stage (Thorlabs, PT1), had a ’manual’ control mode, where the position of the fly was controlled by the experimenter using a joystick, and an ’auto’ control mode, where the fly was moved along a trajectory through previously defined key positions (see section 1.1 for details). First, the head of the fly was positioned (in ’manual’ mode) in the field of view of the microscope with the help of three cameras. Two cameras (Basler acA640-750um) had a large field of view for easily positioning the fly in the center of the microscope, and one camera provided the view through the microscope.

After moving the fly’s head into focus, ’manual’ mode was again used to move the fly’s head to 6 key positions (each with three spatial coordinates), defining the trajectory for laser cutting (see Fig. 1b, center). After defining the key positions, another motorized micromanipulator in ’manual’ mode was used to move a metal shim with a v-shaped incision over the fly’s head to protect the eyes from laser light. The motorized micromanipulator with the fly attached was then set to ’auto’ mode, so that the fly’s head was automatically moved across a trajectory through the previously defined key positions. Finally, the laser shutter was manually opened to cut the cuticle in one pass over the defined key positions. The entire laser surgery process is shown in Supplementary Video 1.

### 1.3 Microrobotic arm

A 3D model of the microrobotic arm is shown in Fig. 2a. Fig. 2c shows the setup with the microrobotic arm with forceps and the dissection microscope. The microrobotic arm was built using translation and rotation stages from Thorlabs that were motorized with stepper motors (Nema 17) (see Fig. 2a), similar to the motorized micromanipulator described in section 1.1. The microrobotic arm had the following joints: three motorized translational joints (Thorlabs PT3), *t*_0_, *t*_1_, and *t*_2_, three rotational joints *r*_0_, *r*_1_ and *r*_2_, and opening and closing of the forceps (Thorlabs MT1), *g*, by actuating one of the forceps arms, as shown Fig. 2b. The rotational joint *r*_0_ was a swivel mount that was used to easily remove the forceps from under the dissection microscope for cleaning the tip of the forceps with a paper tissue during dissections (see Fig. 2d). On the other hand, *r*_1_ and *r*_2_ used a rotational stage (Thorlabs CR1), and only *r*_2_ was motorized with an attached motor. Micro-stepping of the stepper motors (3200 steps per revolution) together with gear reduction, provided a resolution of 0.10*μ*m per step in translational joints, including the joint closing and opening the forceps, *g*, and 0.002 degrees per step in the rotational joint *r*_2_. Two additional one-axis translation stages (Thorlabs MT1) were used without motors to correct the offset between the tip of the forceps and the center of rotation of the *r*_2_ joint to ensure that the rotation occurs around the tip of the forceps. To control the five stepper motors, similar electronics were used to the one in the motorized micromanipulators. TMC2130 drivers were used to move the stepper motors, which were plugged into two CNC shields and these connected to a Teensy board 4.0 (which allows faster communication with the computer and therefore leads to smoother control), which was connected to a host computer through a USB port and received velocity commands for each motor. All the joints of the microrobotic arm were controlled with a single joystick (Logitech Extreme 3D pro joystick) through custom software in Python (host computer) and c++ (on the Teensy board). Velocity sensitivity could be adjusted on the joystick for faster or finer movements in each joint.

### 1.4 Fly stocks and preparation

We used 8-10 days old female flies expressing GCaMP7f in wedge nuerons (UASGCaMP7f;R60D05-GAL4^12^). Flies were reared at 25 degrees with a 12 hour day-night cycle. During long-term imaging, the same time schedule was followed for switching the VR stimulus on and off.

Flies were randomly picked from a vial, briefly anesthetized on ice and attached to a pin mounted on a three-axis micromanipulator with UV glue after immobilizing them in an elongated opening in an aluminum wedge on a cold plate a 4 degrees Celsius^7, 22^. The head of the fly was immobilized by additionally adding two drops of glue with a pin between the thorax and the head close to the eye or occasionally covering the top part of the eye. The pin with the fly was then transferred to the laser cutting setup and an opening was cut into the cuticle as described below (section). The cut cuticle and underlying air sacs were removed with a microrobotic arm with forceps as described below (section). A drop of a glue (DETAX, Freeform, 02204) was placed in the opening and distributed over the edges of the cuticle around the opening with a pin, and then cured with UV light (UV gun as in^7^) for twice 15 seconds from each side of the fly’s head. Flies were then detached from the pin by gently pushing against the thorax close to the attached pin with forceps.

For each experiment we prepared up to six flies. Each surgery took about 20 minutes. After surgery, flies were glued to a microscope cover slide (22 mm × 22 mm, thickness No. 1, Cat. No. 631-0124) while cold anesthetized, either after being transferred to a vial (flies 3 and 4) and left to recover over night, or directly after surgery and left to recover for about 30 minutes (flies 1 and 2).

To attach flies to the cover slide, UV glue (Norland Optical Adhesive 68^9^) was distributed at the center of a cover slide, in a thin layer to minimize optical aberrations. The fly was positioned upside down with its head and part of the thorax on top of the glue layer under a dissection microscope (Leica M205 C, Planapo 10x objective) with forceps, gently pushed against the surface and the glue was cured with UV light for twice 15 seconds from different angles. Additionally, in flies 3 and 4 the antennae were covered with UV glue, to exclude that potential activity in the EB during immobility could be due to movement of the antennae.

For imaging, the microscope slide was fixed to a custom-designed aluminium holder in the microscope using two screws (Fig. S2a). The holder was further attached to a micromanipulator using a magnetic mount (Thorlabs KB25) and the fly was centered on the ball^7^. We then monitored neural activity of the fly with the microscope and its walking behavior on the ball for about 5 minutes and either started the long-term experiment or tested a different fly if imaging quality or behavior weren’t satisfactory. A total of 8 flies were selected for long-term imaging after surgery, 4 of them (all glued to the glass immediately after surgery) died after at least 12 hours of recordings, the other flies lasted between 2 and 7 days (flies 1,2,3 and 4).

**Figure S2.**
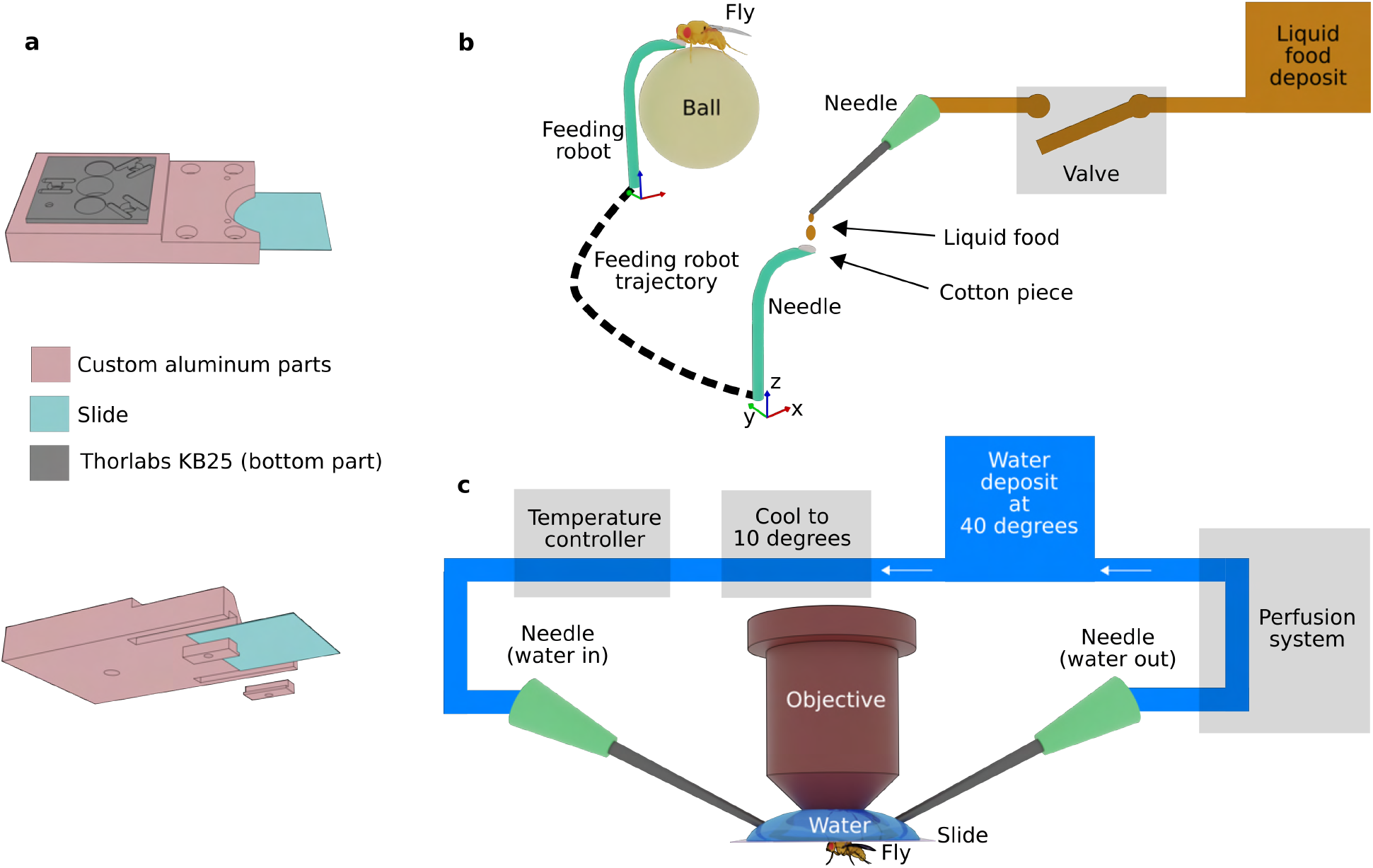
**a** Different views of holder for microscope cover slide with glued fly. **b** Components of feeding system. A needle with a piece of cotton inserted at the tip, is parked in standby under another refilling needle. Refilling is controlled by a valve that opens and closes every 2 minutes, connected to a liquid food deposit. After 4 hours of standby, the feeding needle moves along a trajectory to feed the fly on the ball. **c** Temperature control of the water between the objective and microscope cover slide. Water in a container is heated to 40 degrees and a perfusion system circulates the water to a cooling chamber where it is cooled down to 10 degrees. From there, water is heated again to the desired temperature by a temperature controller and finally circulated under the objective, as commonly done to prevent formation of bubbles under the objective.

Liquid fly food was prepared at the beginning of the experiment and stored at 4 degrees Celsius. 0.5 L of distilled water, 50 g extracted yeast, and 25 g sucrose was heated to 60 degrees for 10 minutes. After cooling down to room temperature, 0.75 g methyl 4-hydroxibenzoate, 1.25 mL Ethanol and 1 mL Propionic acid were added. The food was stored at 4 degrees. The experiment’s liquid food deposit was refilled once a day with a mixture of 200 mL liquid food and 200 mL of distilled water.

Beam profiles were measured with z-stacks recorded with 1*μ*m diameter fluorescent beads (average of 10 stacks). Fig. 3c (semitransparent red and blue lines) shows measured normalized beam profiles fitted with the following Gaussian function:

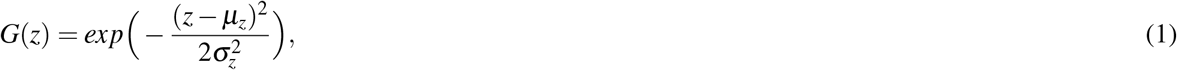

with maximum *μ_z_* and standard deviation 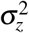 (Fig. 3C, solid red and blue lines).

### 1.5 Temperature control

We used a perfusion system (Multichannel Systems, PPS2 Peristaltic Perfusion System) to control the temperature during imaging. This perfusion system continuously circulated water between the objective and the cover slide in a closed loop at a desired temperature. Water was first heated to 40 degrees in a beaker and then cooled down to 10 degrees inside the tube leading to the objective before reaching the final temperature in a temperature controlled needle dispensing it onto the microscope slide (Mutlichannel Systems, PH01 Perfusion Canula, only heating). Water was removed through a second needle, as shown in Fig. S2c. Preheating of water to 40 degrees before cooling was necessary to avoid the formation of bubbles under the objective. The temperature at the objective was set to 22 degrees for flies 1 and 2, and to 20 degrees for flies 3 and 4.

### 1.6 Virtual Reality setup

Ball motion was tracked with a single camera (Basler acA640-750um) and a single bright stripe was projected onto a five-sided screen using a virtual reality (VR) system based on two DMDs updated at 120 Hz^15^. Ball motion was tracked using optical flow^15^ at 300 Hz. Compared with^15^ ball tracking frame rate was decreased to increase the camera exposure time and reduce the required IR light intensity. The screen was made of semi-transparent paper (Hobbycut Mylarfolie Schablonen-Material) that was attached to a pentagonal frame printed with a 3D printer (Prusa, i3 MK3S) out of transparent PLA filament. A total of four cameras were monitoring the behavior of the fly and the feeder through the semi-transparent screen. The screen had a radius of 30 mm and a height of 70 mm. The fly was at the center of the pentagonal screen, at a height of 20 mm from the top.

A laser (Toptica, ibeam Smart, 488 nm) was used as a light source and was modulated to only switch on during the turnaround of the resonant mirror where imaging data is not recorded^23, 24^. Additionally laser line and complementary emission filters were used to reduce the light on the PMT.

### 1.7 Hardware and software synchronization

Three computers were connected in a local network. The flow of information between the three computers and experiment hardware is shown in Fig. S3. One computer (Windows 10) ran the VR software which was integrated with ROS (Robot Operating System) for Windows, and published through the network in a ROS topic at 300 Hz the time stamp of each frame collected by the camera which tracked the movements of the ball (ball time stamp). This ball time stamp was used as the common time for all events throughout the entire experiment. A second computer (Windows 10) running Scanimage^18^ integrated with ROS (ROS toolbox in Matlab R2016b), was subscribed to the ball time stamp ROS topic. This modified Scanimage version additionally saved a file containing the ball time stamp corresponding to each Scanimage imaging frame. Scanimage ran continuously in its loop mode while being subscribed to a ROS topic to receive grab and stop commands which were sent from a third computer to control the timing of imaging recordings. This third computer (Ubuntu 18.04 LTS and ROS melodic) ran Roscore and other ROS nodes that were also subscribed to the ball time stamp ROS topic. The ROS nodes in this computer were custom-developed in Python and included: a node that controlled the day and night cycle by modulating the power (on and off) of the VR laser beam, and stored the state of the laser with the corresponding ball time stamp; a node to control the robotic feeder, which stored the actions of the robot in the ’auto’ mode with the corresponding ball time stamp; software controlling the timing for opening the liquid food valve; a node showing all frames at 10 Hz, combining different views of the fly (front, zoomed in front, left, zoomed in left, and left top) from five different cameras (Basler Basler acA640-750um), and stored each camera frame (fly view frame) with its corresponding ball time stamp; and software that published in a ROS topic grab and stop commands that were received by Scanimage to record imaging data.

**Figure S3.**
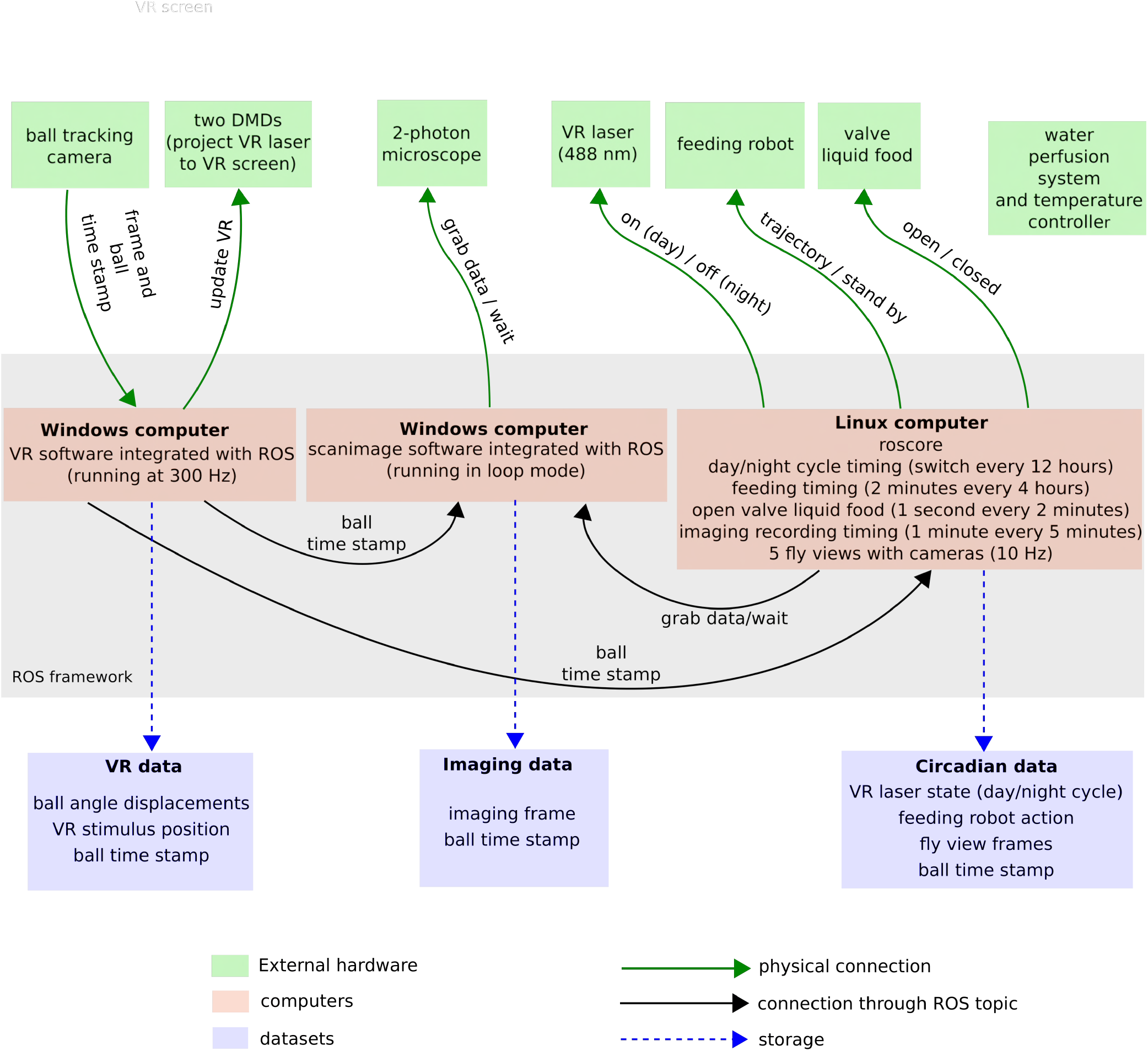
Schematic of setup control. Three different computers communicate trough the ROS framework synchronized to the ball tracking time. The first computer runs the VR software that obtains the ball tracking data and projects the bright stripe onto the VR screen using two DMDs, similar to^15^. The second computer controls the 2-photon microscope and receives commands for recording imaging data. The third computer controls the day and night cycle, the feeding robot, and obtains views of the fly from different cameras. Each computer stores a dataset for analysis.

### 1.8 Data storage and preprocessing

Each computer stored a dataset, resulting in three datasets per experiment that included VR data, imaging data and circadian data, as shown in Supplementary Fig. S3. The three datasets were joined together and kept in a external storage unit for data analysis. The amount of data for single flies ranged from 0.5 Tb up to to 1.5 Tb. Scanimage time stamps were synchronized with ball time stamps with a conversion factor measured form simultaneously recorded data, and either time stamps were used.

For data analysis, we described the data of a single fly in DataFrames using the library Pandas, which contained, for example, the link to the file directory of each single imaging frame, together with its corresponding ball time stamp, the corresponding interpolated VR (ball displacements and VR stimulus position), circadian data (light state and feeding robot actions), and the corresponding fly view frame closest to that particular time stamp. This way of organizing the data allowed access to all data without having to load them into memory. We divided the data into continuous recordings of imaging data (trials) lasting for 60 seconds in experiment 1 and 2 and 30 seconds for experiments 3 and 4.

### 1.9 Imaging data analysis from EPG neurons

The first step for data analysis of each trial was to correct the lateral motion between two-photon images (referred as *xy* motion). First, for a trial a template was constructed by averaging over the first 20 frames. The *xy* offset for the remaining frames in the trial with respect to the template was computed using a phase correlation algorithm implemented in the Skimage library^25^. Using the *xy* offset, all frames in the trial were aligned, which included 7200 (for flies 1 and 2) or 3600 frames (for flies 3 and 4), and the average of these frames was used as a common template for all the remaining trials (see Fig. S4a). The offset of every frame in all trials was computed with respect to the common template using the same algorithm. To improve the *xy* motion correction, we averaged up to 20 frames and used either a Gaussian or a median filter to smooth frames, as shown in Fig. S4a. These setting were selected once for each fly and maintained over all trials.

**Figure S4.**
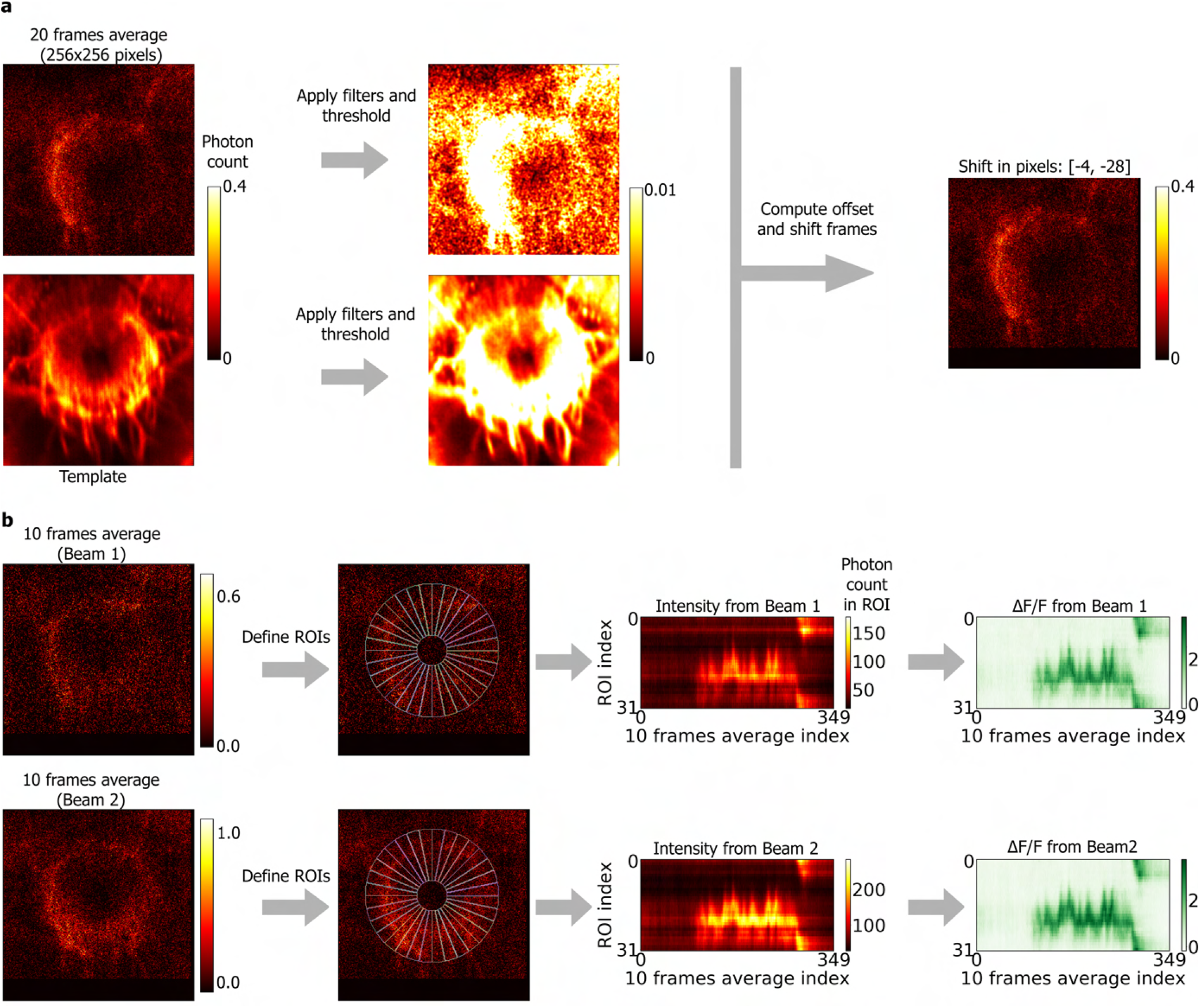
Analysis of imaging data. **a** In each trial, lateral motion is corrected using a template. Left side: 20 frames are averaged and Gaussian and threshold filters are applied to both the template and the 20 averaged frames (center). A phase correlation algorithm is used to compute the offset from the template and the 20 averaged frames are shifted to match the template (right). **b** 10 frames were averaged in the planes recorded from each beam (left) and 32 ROIs (center left) were defined to calculate the intensity over time within each ROI (center right). The change in fluorescence Δ*F/F* was calculated for each beam (right) and the final change in fluorescence was the average of the two.

After correcting lateral motion, a total of 32 wedge-shaped regions of interest (ROIs) were defined and the sum of intensity in each ROI was computed for each frame. For each trial, two intensity matrices, *I*_1_ and *I*_2_, corresponding to the two simultaneously recorded imaging planes were obtained. Each intensity matrix included the intensity in each ROI computed from 10 averaged frames, providing a time resolution of 0.17 seconds. Both matrices had a size of 32 × *T/m*, where *T* is the number of frames in a trial per simultaneous plane, and *m* = 10 is the number of averaged frames (Fig. S4b). We then computed the change in fluorescence for each recorded plane, (Δ*F/F*)_1_, and (Δ*F/F*)_2_, as

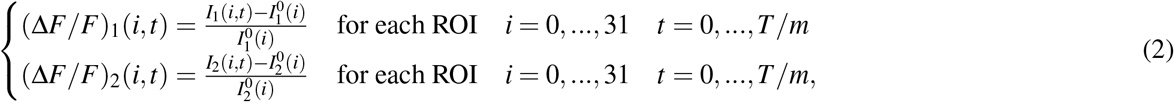

where 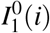 and 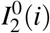 are the baseline activity of each ROI *i* for each plane defined as the mean of 10% of imaging frames with lowest activity in ROI *i*. We finally combined the change in fluorescence in each plane to obtain the fluorescence matrix:

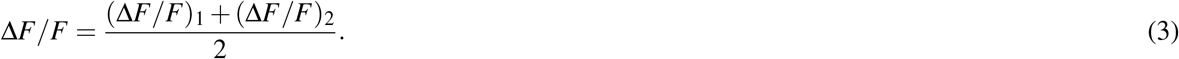

With this fluorescence matrix, we computed the amplitude of the bump of activity in wedge neurons, *A*(*t*), as the maximum value in Δ*F/F* at a given time *t*:

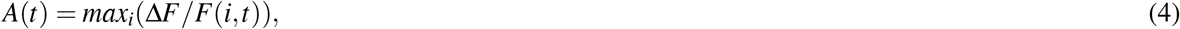

and the full width at half maximum, FWHM(t).

To observe how the bump amplitude and bump width changed over time, we computed the mean bump amplitude in the trial, < *A* >, and the mean bump full width at half maximum in each trial, < FWHM >,

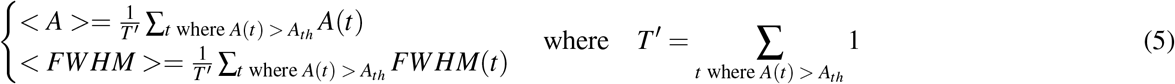

filtering out times where the bump amplitude, *A*(*t*), was lower than a threshold, *A_th_* = 1. This avoided averaging over times where a bump was not visible when the fly was not moving. Figures 4, S5, S6, S7 and S8 show the mean bump amplitude in each trial, < *A* > (fourth row). The fifth row shows < *FWHM* >.

We then computed the bump position in wedge neurons in each trial using algorithm 1. For this, first the time *t* where the change in fluorescence is largest per trial, *t_max_*, was determined. Then the population vector average (*PVA*) at time *t_max_* was calculated as *PVA*(*t_max_*) = *pva*(*t_max_*), with the function *pva*(*t*) for time *t* defined as:

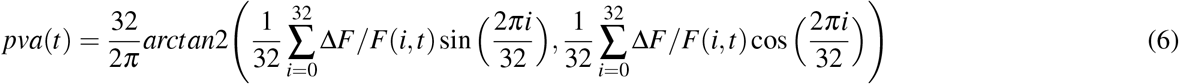

Finally, the *PVA* was computed backwards and forwards in time from *t_max_*. If the maximum change in fluorescence at time *t* was larger than the threshold *F_th_*, the *PVA* was computed as *PVA*(*t*) = *pva*(*t*). Otherwise it was assigned the next or previous value, *PVA*(*t* + 1) or *PVA*(*t* − 1), depending on the direction of time:

#### Algorithm 1

Calculation of *PVA* for each trial

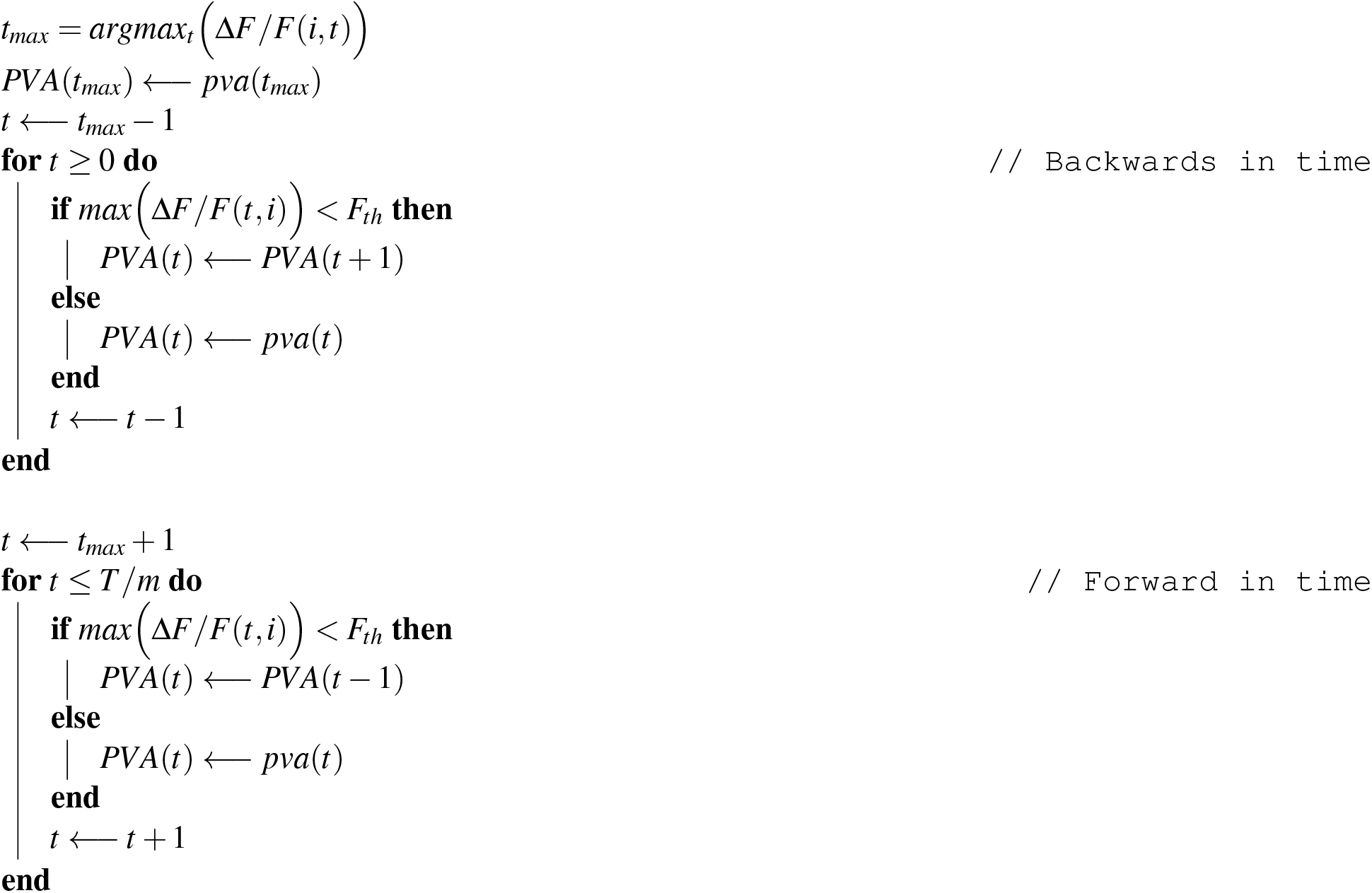

The PVA in the range of [0, 32] was scaled to [− 180, 180], corresponding to the VR orientation, *R*(*t*). The PVA and VR orientation were both unwrapped to compute the PVA-VR orientation offset, *PVA*_0_, for each trial. The PVA-VR orientation offset was computed as the mean of the offset between the two variables at times when the bump amplitude, *A*(*t*), was larger than the threshold *A_th_*:

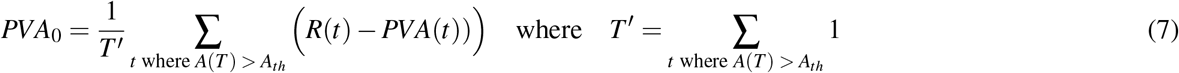

Then, the *L*_1_ error between the PVA, with offset *PVA*_0_ subtracted, and the VR orientation was found as:

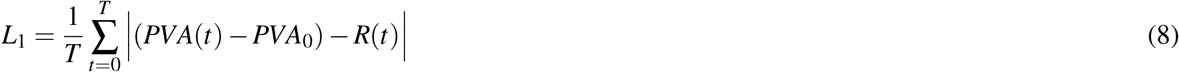

This error indicated how well the PVA encoded the VR orientation. In trials where the fly was not moving, this error was close to zero. Values of *L*_1_ error in trials where the mean velocity during the trial *v^trial^* was lower than a threshold *v^trial^* < *v_th_* = 0.1*mm* could therefore be removed. The values for the *L*_1_ error are shown in the 7th row of Fig. 4, and Supplementary Figures S5, S6, S7 and S8.

**Figure S5.**
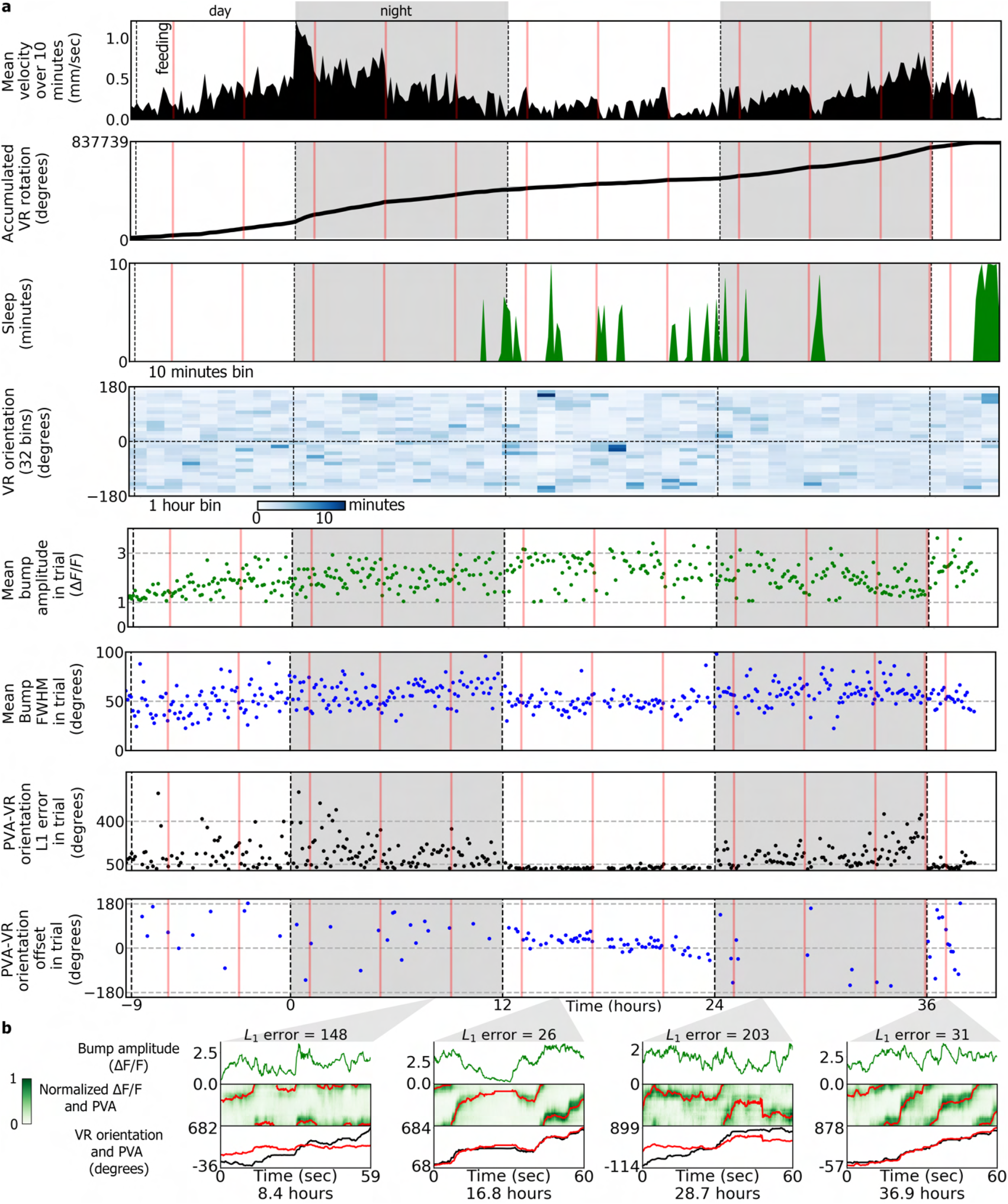
Imaging and behavior experiment for fly 2. See figure legend of Fig. 4 for details.

**Figure S6.**
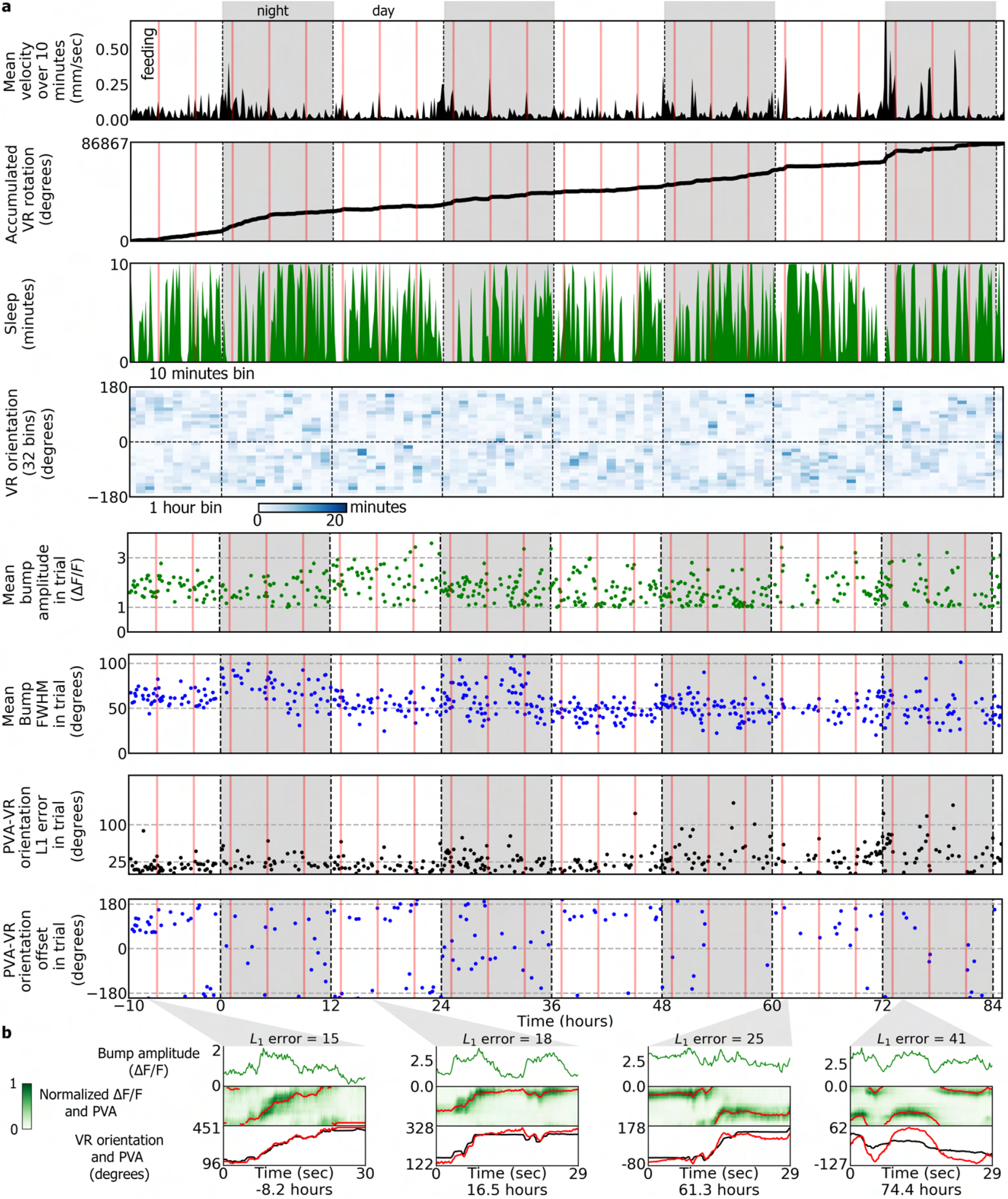
Imaging and behavior experiment for fly 3, first part. See figure legend of Fig. 4 for details.

**Figure S7.**
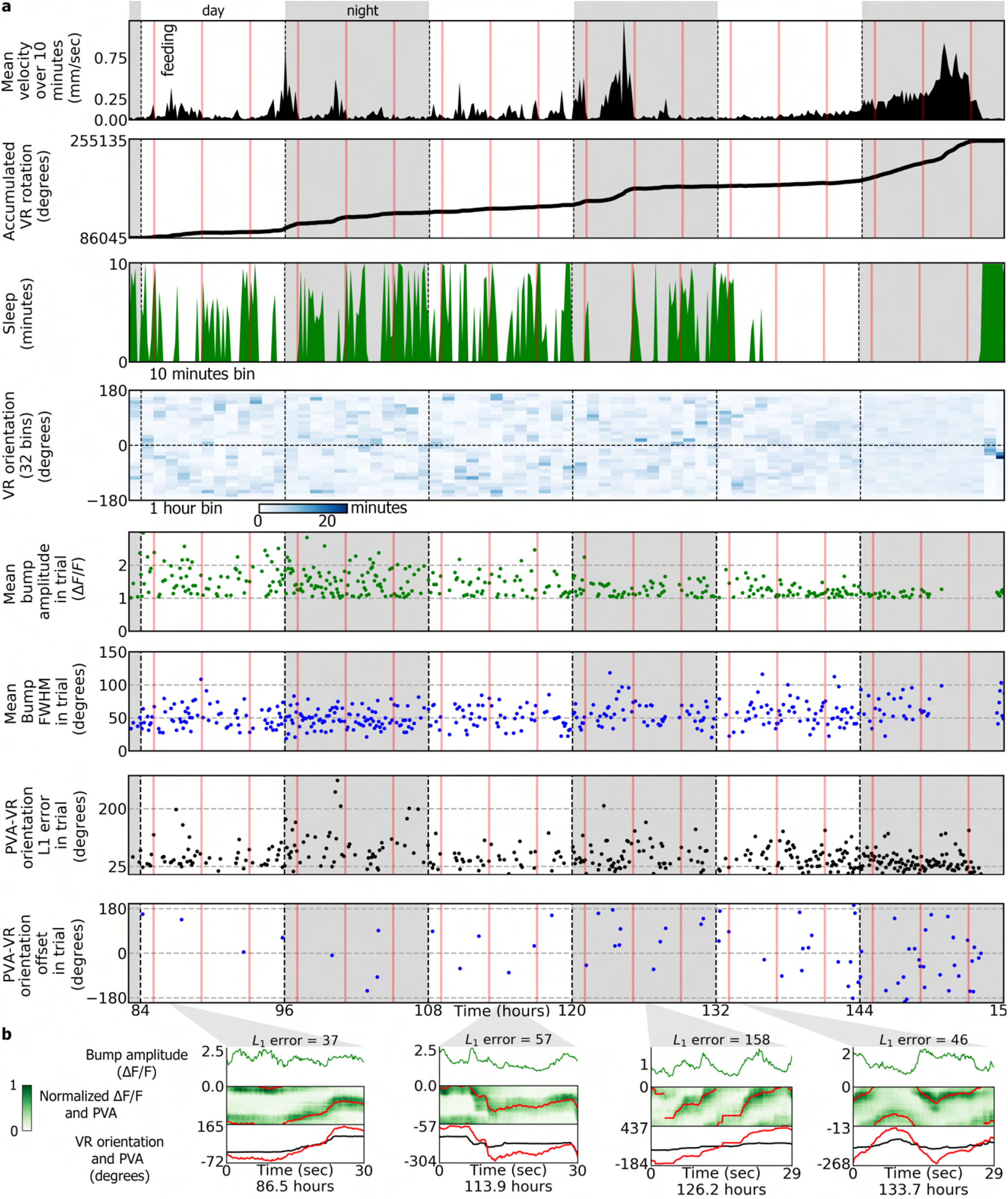
Imaging and behavior experiment for fly 3, second part. See figure legend of Fig. 4 for details.

**Figure S8.**
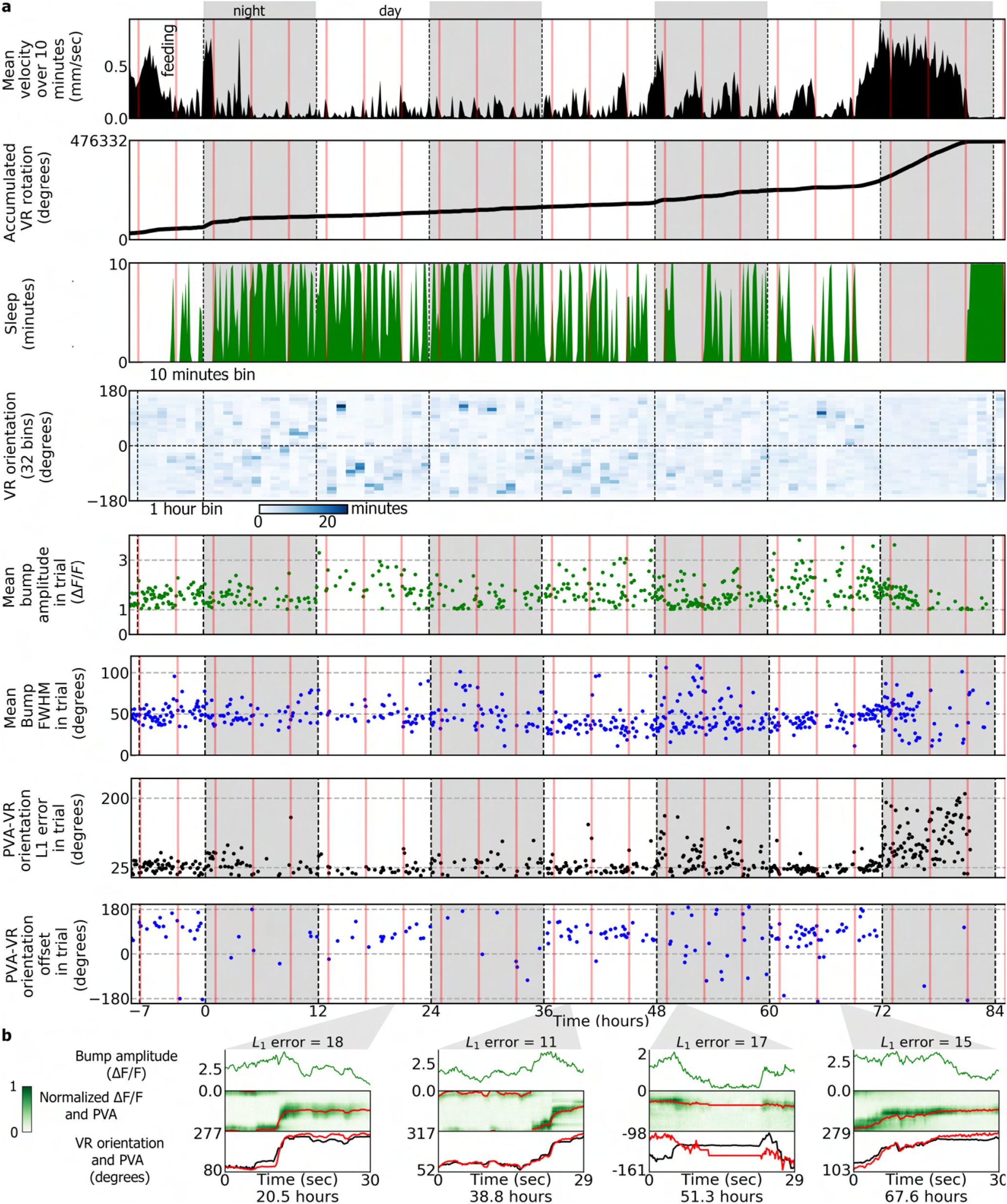
Imaging and behavior experiment for fly 4. See figure legend of Fig. 4 for details.

**Figure S9.**
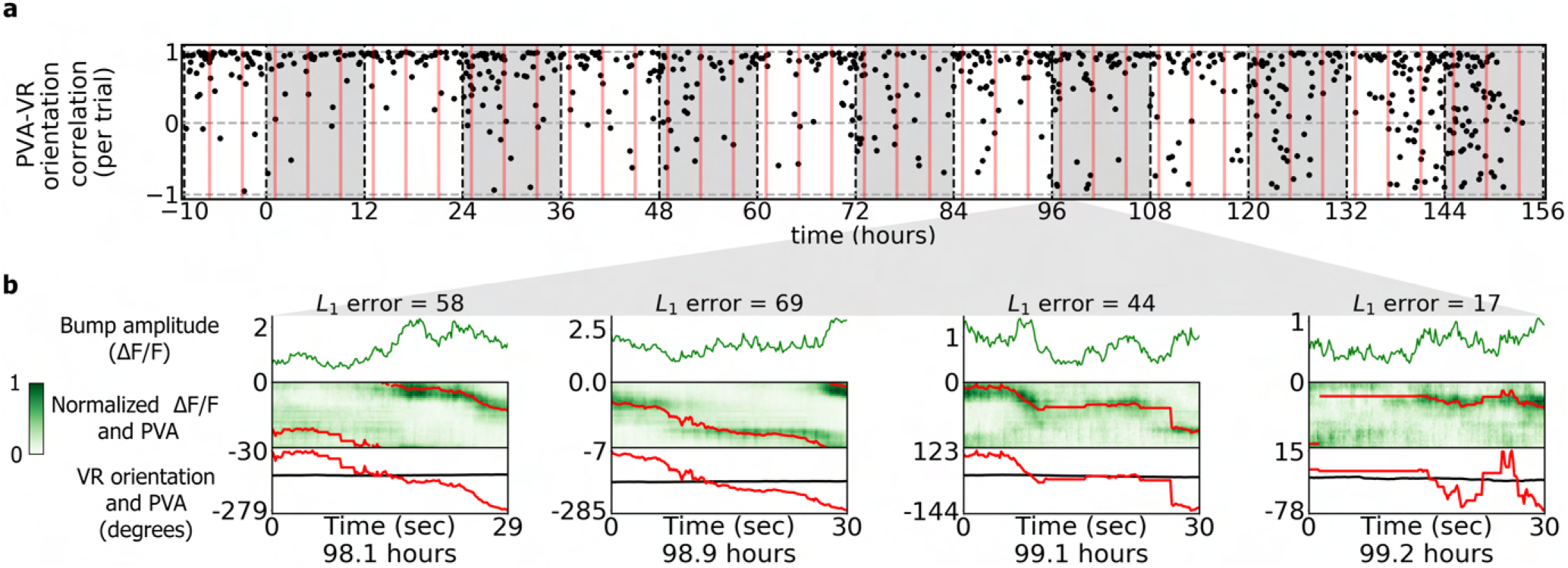
PVA-VR correlation and bump drift in fly 3. **a** Pearson correlation between the PVA and VR orientation in each trial remains high for all trials, whereas the L_1_ error increases after the 5th day (Fig. S7, 7th row). **b** Examples of trials during the night where the bump drifts while the fly is standing still on the ball. The antennae of the fly were glued. Such strong autonomous dynamics was only observed after the L_1_ error increased during the day, potentially due to phototoxicity.

**Figure S10.**
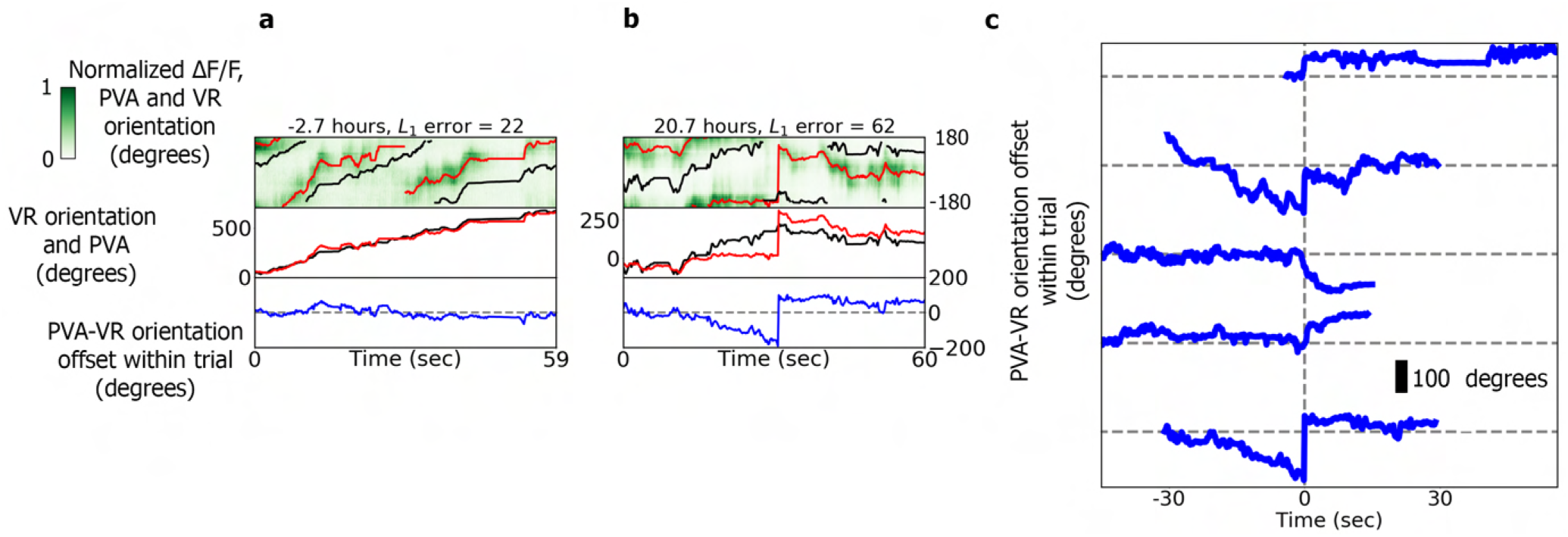
Delays in bump position updating in fly 1. **a** Example of a trial where the bump is moving as expected during the first day. **b** Example of a trial where the bump doesn’t accurately track the stimulus during the second day and is delayed in one position before moving on. **c** PVA-VR orientation offset in 5 trials where the same phenomenon was observed during the second day. No delays were observed during the following day.

In addition, to analyze changes in PVA-VR orientation offset, *PVA*_0_, over the course of the experiment, we selected the offset in trials where both the mean velocity in the trial was higher than the threshold *v_th_*, and where the *L*_1_ error was lower than a second threshold, 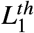. This threshold was selected at 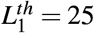 degrees for flies 3 and 4, and 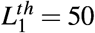 degrees for flies 1 and 2, since the trial duration for the latter was twice as long and therefore larger errors accumulated. PVA-VR orientation offsets with high *L*_1_ error were removed since in this case the *PVA*_0_ offset was not reliable. Cases where the *L*_1_ error was large included trials during the night (See Fig. 5). Another source of large *L*_1_ error were fast rotations of the fly that resulted in very fast bump movement with low fluorescence intensity that was therefore not recognized by the algorithm 1. All parameter values for analysis of each fly are shown in Supplementary Table S2.

**Table S1.**
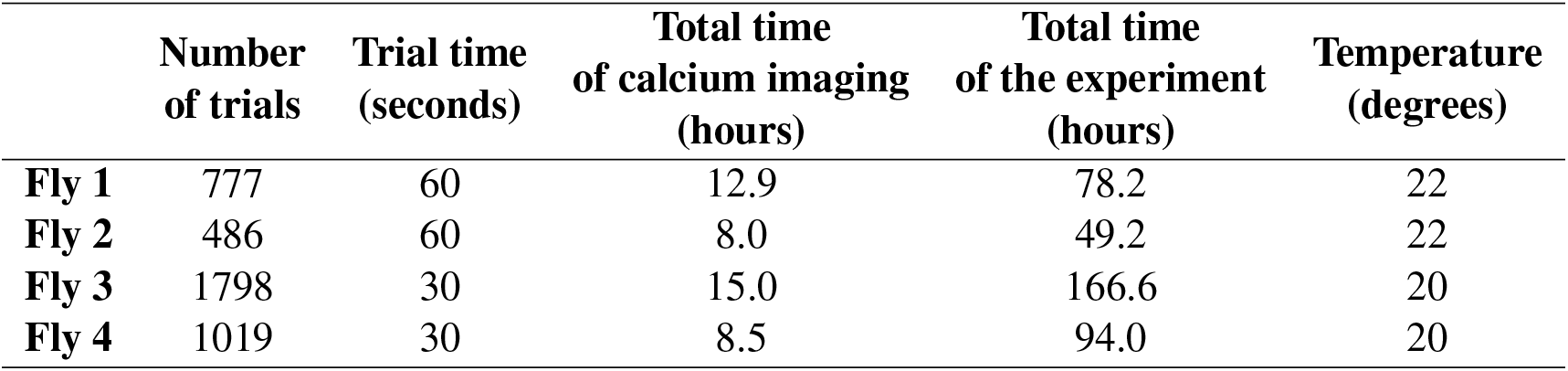
Trials, timings and temperatures of the long-term imaging experiments for each recorded fly.

|  | Number<br>of trials | Trial time<br>(seconds) | Total time<br>of calcium imaging<br>(hours) | Total time<br>of the experiment<br>(hours) | Temperature<br>(degrees) |
| --- | --- | --- | --- | --- | --- |
| **Fly 1** | 777 | 60 | 12.9 | 78.2 | 22 |
| **Fly 2** | 486 | 60 | 8.0 | 49.2 | 22 |
| **Fly 3** | 1798 | 30 | 15.0 | 166.6 | 20 |
| **Fly 4** | 1019 | 30 | 8.5 | 94.0 | 20 |

**Table S2.** Parameter values used in each experiment for data analysis.

| | Averaged<br>imaging frames<br>for lateral<br>motion correction | Averaged<br>imaging<br>frames<br>in template | Amplitude<br>threshold<br>$A_{th}$<br>($\Delta F/F$) | Fluorescence<br>threshold<br>$F_{th}$<br>($\Delta F/F$) | Linear velocity<br>threshold<br>$v_{th}$<br>(mm/sec) | $L_1$<br>error<br>threshold<br>(degrees) | Time range<br>for statistical<br>analysis<br>(hours) |
| --- | --- | --- | --- | --- | --- | --- | --- |
| Fly 1 | 20 | 7200 | 1 | 0.4 | 0.1 | 50 | [-3,70] |
| Fly 2 | 20 | 7200 | 1 | 0.6 | 0.1 | 50 | [0, 39] |
| Fly 3 | 10 | 3600 | 1 | 0.6 | 0.1 | 25 | [-10, 84] |
| Fly 4 | 10 | 3600 | 1 | 0.6 | 0.1 | 25 | [-7, 68] |

### 1.10 Behavioral data analysis for each fly

Fly behavior was extracted from ball movements recorded over the entire experiment, also in between imaging trials. Ball tracking^15^ provided the angular displacement of the ball along the three axes of rotation, (*b_x_*(*k*), *b_y_*(*k*), *b_z_*(*k*)), at a rate of 300 Hz for time stamp *k*. We first computed the linear velocity of the fly in mm/sec as:

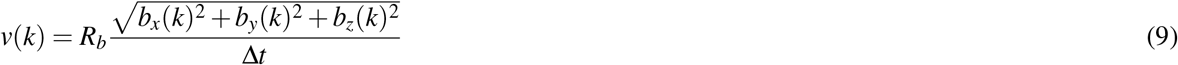

where *R_b_* = 3 mm is the radius of the ball and Δ*t* = 1/300 sec is the time between consecutive frames. We then computed the mean velocity of the fly over 10 minutes by averaging v(k) over 10 minute bins (Fig. 4 and Supplementary Figures S5, S6, S7 and S8, first row).

Since the ball tracking algorithm shows slow drift due to imaging noise, we defined a velocity threshold, *v_th_* = 0.1 mm that was used to detect sleep events. Sleep was defined as the velocity being below *v_th_* for at least 5 minutes^1^. To display sleep events, we finally computed the amount of sleeping time in bins of 10 minutes (Fig. 4 and Supplementary Figures S5, S6, S7 and S8, third row). That the fly was not moving during these times could additionally be verified by simultaneously recorded videos of the fly on the ball.

Walking direction of the fly was computed from the accumulated VR rotation (Fig. 4 and Supplementary Figures S5, S6, S7 and S8, second row). The distribution of VR orientation over time in a histogram with a bin width of VR angular range [−180, 180] divided by 32 (wedges) over 1 hour is shown in Fig. 4 and Supplementary Figures S5, S6, S7 and S8, fourth row.

### 1.11 Statistical analysis of behavior and imaging data

For each fly we compared the following quantities across trials between day and night: (i) walking velocity, (ii) mean bump amplitude, (iii) mean bump FWMH, and (iv) L_1_ error. Walking (or ball) velocity was averaged over 10 minutes, as described in section 1.10, while the other quantities were obtained for each trial, as explained in section 1.9. For (ii), (iii) and (iv) only trials where the fly was moving were included (only trials with a mean velocity larger than a threshold velocity *v_th_*). Statistical significance was assessed with Kolmogorow-Smirnow (KS) test (in Scipy^26^). Additionally, only time intervals where wedge neurons tracked the bright stripe in the VR well were included (see Table S2, last column). All comparisons were statistically significant (p < 0.05), as shown in Fig. 5 i), ii), iii) and iv).

In addition, the effect of feeding on (v) the velocity of flies, (vi) the mean bump amplitude per trial and (vii) the bump FWHM per trial was analyzed. For comparison, 1 hour before feeding and 1 hour after feeding, both during the day and during the night, and within the same time interval, as in the previous analysis (see Table S2), were considered. For mean velocity values over 10 minutes, we performed two t-tests, one with values during the day and the other with values during the night. Statistical significance was found for 3 out of 4 flies during the night, while only fly 1 showed statistical significance during the day, as shown in Fig. 5 v. For quantities (vi) and (vii), again only epochs with velocity larger than *v_th_* were considered. Finally, two t-tests were performed for each quantity (vi) and (vii) during both the day and night (see Fig 5 vi and vii)

